# Phase-Amplitude Coupling of Neural Oscillations can be Effectively Probed with Concurrent TMS-EEG

**DOI:** 10.1101/273482

**Authors:** Sarah Glim, Yuka O. Okazaki, Yumi Nakagawa, Yuji Mizuno, Takashi Hanakawa, Keiichi Kitajo

## Abstract

Despite the widespread use of transcranial magnetic stimulation (TMS), knowledge of its neurophysiological mode of action is still incomplete. Recently, TMS has been proposed to synchronise neural oscillators, and to thereby increase the detectability of corresponding oscillations at the population level. As oscillations in the human brain are known to interact within nested hierarchies via phase-amplitude coupling, TMS might also be able to increase the macroscopic detectability of such coupling. In a concurrent TMS-electroencephalography study, we therefore examined the technique’s influence on theta-gamma, alpha-gamma and beta-gamma phase-amplitude coupling by delivering single-pulse TMS (sTMS) and repetitive TMS (rTMS) over the left motor cortex and right visual cortex of healthy participants. The rTMS pulse trains were of 5 Hz, 11 Hz and 23 Hz for the three coupling variations, respectively. Relative to sham stimulation, all conditions showed transient but significant increases in phase-amplitude coupling at the stimulation site. In addition, we observed enhanced coupling over various other cortical sites, with a more extensive propagation during rTMS than during sTMS. By indicating that scalp-recorded phase-amplitude coupling can be effectively probed with TMS, these findings open the door to the technique’s application in manipulative dissections of such coupling during human cognition and behaviour in healthy and pathological conditions.

## 1. Introduction

Due to its extensive effects on human perception, cognition and action, transcranial magnetic stimulation (TMS) is nowadays widely used in both basic neuroscientific research (e.g., during investigations of visual awareness [1], attention [2], speech [3] and motor processing [4]) and in clinical practice (with potential treatment domains [see guidelines on therapeutic use [5]] including medication-resistant major depressive disorder [6], post-stroke motor impairment [7], aphasia [8] and schizophrenia [9]). Despite this broad scope of application, knowledge of the precise neurophysiological effects of TMS is still incomplete.

Over the past decade, interest has arisen in the effects of TMS on macroscopic neural oscillations, as measured with non-invasive recording techniques such as electroencephalography (EEG) [10–14]. In this context, Kawasaki and colleagues [13] demonstrated a direct modulation of the temporal dynamics of these oscillations by showing that the consistency of oscillatory phases across stimulation trials, so-called phase locking, is transiently enhanced after single-pulse TMS (sTMS). Even though this effect can occur within a wide oscillatory spectrum, sTMS is assumed to act on intrinsic neural systems, and thus to be most effective for those frequencies that arise naturally within particular cortico-thalamic modules [15]. Accordingly, a highly probable candidate mechanism behind the observed increase in macroscopic across-trial phase locking is the phase resetting of underlying intrinsic oscillators (but see Sauseng and colleagues [16] for a critical discussion of phase locking). Considering that such resets would simultaneously pertain to a multitude of coexistent oscillators, transiently enhanced synchronisation would also unfold within stimulation trials. As Thut and colleagues [12, 17] argued, rhythmic stimulation via repetitive TMS (rTMS) can foster such a synchronisation through neural entrainment, during which individual oscillators start to cycle with the same period as pulses delivered at their eigenfrequency, and thus become more and more aligned to such pulses, and consequently also to each other. Interestingly, this synchronisation or alignment of coexistent neural oscillators has been argued to prevent population-level signal nullifications, and to thereby enhance the detectability of macroscopic oscillations with scalp-based measurement techniques [17]. Associated EEG-recorded oscillatory power increases have de facto been reported for both sTMS [15, 18] and rTMS [12].

To fully appreciate the neurophysiological effects of TMS, it is necessary to consider that the human brain is unlikely to be a composition of neatly separated neural modules whose oscillatory signatures can be manipulated independently from each other. Rather, its essence lies in a myriad of dynamic neural interactions that serve the integration of information across various temporal and spatial processing scales [19]. One promising mechanism for how such integration may be implemented in the brain is through a nested hierarchy of neural oscillations [20]. In particular, studies have shown that the phase of oscillations arising from slower global computations can flexibly modulate the amplitude of faster local oscillations [21–25], a mechanism that might enable the coordination of multiple specialised processing nodes across large-scale brain networks. The functional relevance of such phase-amplitude coupling is supported by findings associating its strength with behavioural outcomes, e.g., success in a visual motion discrimination task [26]. Given that phase-amplitude coupling is an inherent property of neural systems, the alignment of oscillators by TMS should enhance not only the detectability of individual macroscopic oscillations, but also the detectability of their coupling to other oscillations. As this feature would greatly facilitate the investigation of phase-amplitude coupling with non-invasive measurement techniques such as scalp EEG, which often require extensive recordings to cope with only moderate signal-to-noise ratios, its clear demonstration would be of high relevance for both TMS methodologists and cognitive neuroscientists.

Attempts have already been made to demonstrate an enhancement of EEG-recorded phase-amplitude coupling by TMS [27] and other non-invasive brain stimulation techniques, specifically transcranial alternating current stimulation (tACS) [28]. Even though Noda and colleagues [27] demonstrated the enhancement of pre-existing intrinsic theta-gamma coupling within an offline paradigm following several sessions of rTMS in patients with depression, conclusive evidence from a sham-controlled examination of online EEG recordings during TMS in the healthy population is still missing. With the present study, we set out to provide such evidence, thereby using TMS to shed light on the transient modulation of the human brain’s nested oscillations. To this end, we delivered both sTMS and rTMS over the left motor cortex and right visual cortex of healthy participants while simultaneously collecting EEG. To ensure coverage of a wide range of the oscillatory nesting observable in neural systems [20, 29, 30], the enhancement of phase-amplitude coupling relative to sham stimulation was assessed separately for theta-gamma, alpha-gamma and beta-gamma coupling, with the alpha and beta bands, in particular, having been related to the stimulated visual and motor cortex, respectively [15]. The rTMS frequency always equalled the frequency of the slower modulating oscillation to allow for this oscillation’s direct entrainment. We designed the experiments to evaluate the following theoretical reasoning. As enhanced oscillatory power has been reported for both sTMS [15, 18] and rTMS [12], scalp-recorded phase-amplitude coupling should likewise be transiently enhanced for both stimulation paradigms. As both paradigms were further shown to modulate phase dynamics not only locally at the stimulation site, but also with network-wide signal propagation [13, 14], the enhancement of phase-amplitude coupling might likewise propagate across the cortex. Finally, we directly compared the neurophysiological effects of sTMS and rTMS by examining whether an rTMS-induced entrainment of neural oscillators can induce a locally stronger and/or globally more widespread enhancement of phase-amplitude coupling relative to sTMS.

## 2. Materials and Methods

### 2.1. Participants

Fourteen right-handed healthy participants (two females, twelve males; mean age ± SD, 30.8 ± 5.5 years) were recruited in this study. Written informed consent was obtained from all participants prior to the experimental sessions. The study was approved by the RIKEN Ethics Committee and was conducted in accordance with the code of ethics of the World Medical Association for research involving humans (Declaration of Helsinki).

### 2.2. TMS Design

TMS pulses were delivered through a figure-of-eight coil with a 70 mm wing diameter, connected to a biphasic magnetic stimulator unit (Magstim Rapid, The Magstim Company Ltd, UK). Stimulation intensity was fixed at 90% of a participant’s active motor threshold, which was determined for the right first dorsal interosseous (FDI) muscle. During the entire experimental procedure, participants fixated on a central grey cross on a black computer monitor background and wore earplugs to reduce stimulation-evoked auditory potentials in neural activity.

An overview of the experimental design is presented in Figure 1. Each participant received stimulation at three different sites in randomly ordered sessions. In one session, TMS was applied over the left motor cortex (approximately between electrodes C1 and C3, with the exact position being determined by the individual hotspot of the right FDI muscle stimulation; coil handle perpendicular to the central sulcus; antero-posterior current direction of the waveform’s first phase) and in a second session, it was applied over the right visual cortex (between electrodes Oz and O2; coil handle perpendicular to the midsagittal plane). In a third session, sham stimulation was delivered at a location 10 cm above the vertex of the head (electrode Cz; coil handle directed posteriorly). Each of these sessions comprised four different blocks, with each block consisting of 30 trials with inter-trial intervals of 10 s ± 15%. Depending on the block, trials contained either single TMS pulses or trains of five consecutive pulses delivered at 5 Hz, 11 Hz or 23 Hz. The different rTMS frequencies were selected so that they are not multiples of each other.

**Figure 1:**
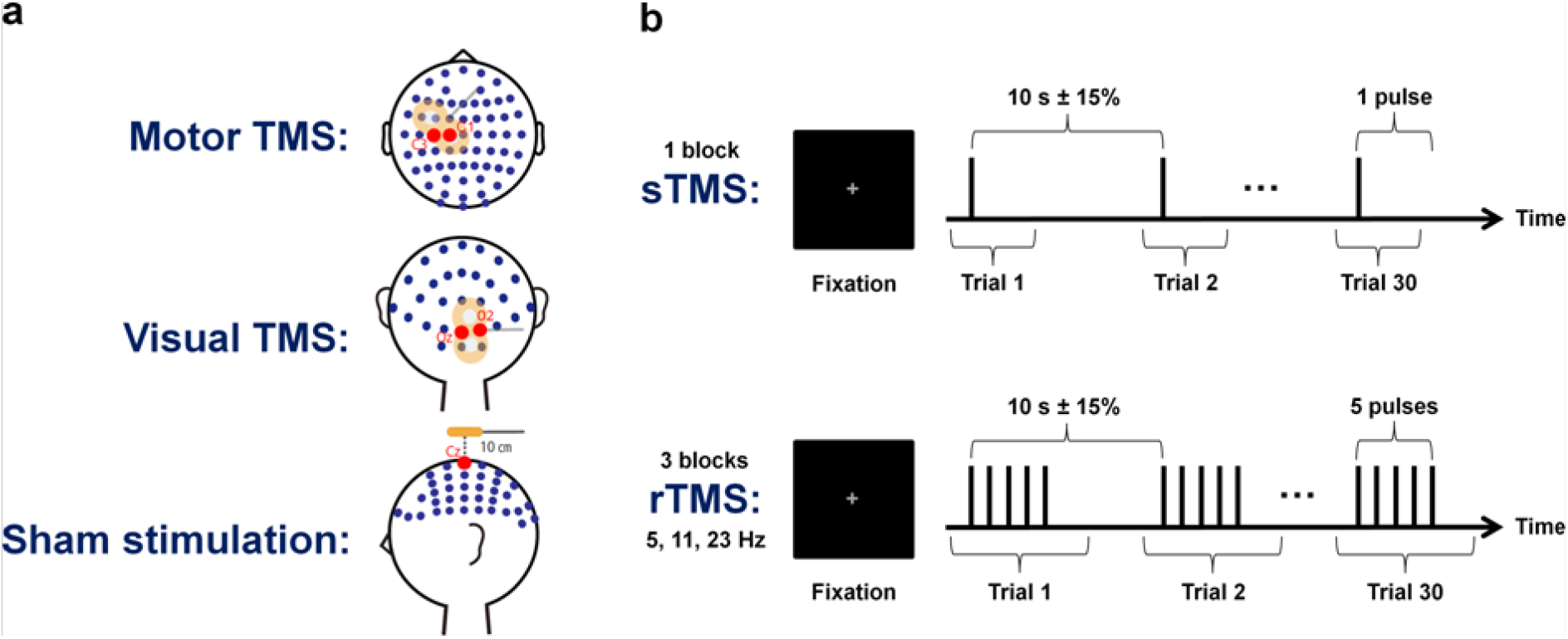
Experimental design. (a) The EEG electrode layout used in the present study is displayed along with the different stimulation sites. In separate sessions, TMS was applied over the left motor cortex (first row), over the right visual cortex (second row) and as sham stimulation 10 cm above the vertex of the head (third row). (b) Each session contained four blocks of 30 trials each, in which we performed sTMS (first row) as well as 5 Hz, 11 Hz and 23 Hz rTMS (second row). During rTMS trials, stimulation was delivered in trains of five consecutive pulses.

### 2.3. EEG Recording and Preprocessing

During the entire stimulation procedure, EEG (left earlobe reference; AFz as ground) was recorded from 63 TMS-compatible Ag/AgCl scalp electrodes (EASY CAP, EASYCAP GmbH, Germany; see Figure 1a for the electrode layout), which were positioned according to the international 10/10 system with lead wires rearranged orthogonally to the TMS coil handle to reduce TMS-induced artefacts [31]. In addition, horizontal and vertical electrooculography (EOG; ground electrode on the left mastoid) was recorded to monitor eye movements and blinks. All signals were sampled at a rate of 5,000 Hz, filtered online from DC to 1,000 Hz and amplified using the TMS-compatible BrainAmp MR plus system (Brain Products GmbH, Germany). Impedances were kept below 10 kΩ.

We preprocessed the EEG data by first segmenting it into epochs starting 2 s before the first (or single) TMS pulse and ending 3 s after the last (or single) pulse of a train, and then re-referencing these epochs to the averaged recordings from electrodes positioned on the left and right earlobe. To remove the TMS-induced ringing artefact in the EEG signals, we substituted all values within an interval of 0–8 ms after each pulse with replacement values estimated using linear interpolation. In those cases where the interval was deemed to be too short via visual inspection, it was manually extended to 12 ms after the pulse. The longer-lasting exponential decay artefact was attenuated by identifying components capturing this artefact with an independent component analysis (ICA), and then removing them from the data [18, 32]. Next, we rejected trials with signal values exceeding ± 200 μV within an interval of −1 s to +1 s around the stimulation to exclude any remaining artefacts. Out of 30 collected trials per block, 24.8 ± 2.6 trials (mean ± SD) were retained. After performing a current source density (CSD) transformation of the surface voltage distribution using spherical splines to reduce the effects of volume conduction [33, 34], the data were down-sampled to a rate of 1,000 Hz.

### 2.4. EEG Analysis

To compute phase-amplitude coupling, we first convolved the preprocessed time series with complex Morlet wavelets *w*(*t,f*) [35, 36]:

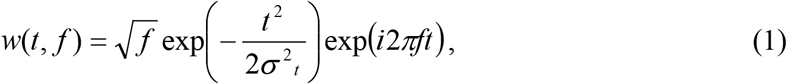

with *t* denoting time, *f* denoting the central frequency of interest, *o’_t_* denoting the SD of the Gaussian window and the number of wavelet cycles within a 6*o’_t_* interval *n_co_* = 3 determining the approximate width of the frequency bands [37]:

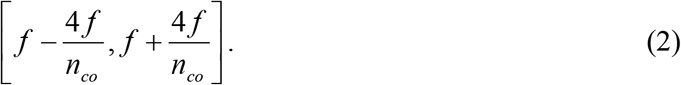

The central frequencies *f* were chosen to be 5 Hz, 11 Hz and 23 Hz for phase extraction and 30 Hz to 45 Hz in 1 Hz steps for amplitude extraction. The upper limit was fixed at 45 Hz to diminish potential artefacts from muscular activity and power line noise. The instantaneous phase *ɸ* at each time point was then defined as the angle of the resulting complex-plane vector with respect to the positive real axis, while the magnitude of this vector was utilised as a measure of instantaneous amplitude *a*. For each combination of phase and amplitude frequency and for each trial time point, we separately computed the event-related phase-amplitude coupling (ERPAC) *ρ_ɸa_*, which was defined as the circular-linear correlation of phase and amplitude values across stimulation trials [38, 39]:

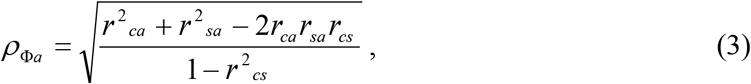

where *r_ca_* = *c*(cos*ɸ*[*n*], *a*[*n*]), *r_sa_* = *c*(sin*ɸ*[*n*], *a*[*n*]) and *r_cs_* = *c*(sin*ɸ*[*n*], cos*ɸ*[*n*]) with *c*(*x, y*) being the Pearson correlation between *x* and *y*.

As the sixteen amplitude frequencies were pooled together in each ERPAC analysis, three different phase-amplitude combinations existed (5 Hz, 11 Hz or 23 Hz phase coupled to amplitudes at 30–45 Hz, i.e., theta-gamma coupling, alpha-gamma coupling and beta-gamma coupling), which were examined separately for motor and visual TMS. Analyses of the resulting six conditions focused on contrasting motor or visual TMS with sham stimulation to account for any indirect effects of stimulation and were performed either individually for sTMS and rTMS (first and third analysis) or directly compared the two stimulation paradigms (second and fourth analysis), as detailed below. Whenever statistical tests were performed, the (multiple-comparison-corrected) significance level was set at *p* ≤ .05. With regard to rTMS, it is important to note that the frequency of a condition’s respective phase angle time series always corresponded to the applied stimulation frequency. This approach allowed us to directly assess how targeting a particular oscillation via repetitive stimulation affected this oscillation’s scalp-recorded coupling to faster oscillations.

We first examined whether sTMS and rTMS led to an increase in phase-amplitude coupling at the stimulation site by analysing ERPAC as a function of amplitude frequency and time, spanning −0.5 cycles to +4.5 cycles of a condition’s phase-providing oscillation around the onset of the first (or single) pulse. Statistically significant enhancements of ERPAC were determined via nonparametric permutation testing in the following way. To evaluate the observed set of time-frequency representations encompassing the ERPAC data from the two local electrodes of interest (C1 and C3 for motor TMS; Oz and O2 for visual TMS), the two modes of stimulation (TMS and sham) and each of the fourteen participants, we created 500 sets of corresponding surrogate representations by computing ERPAC between the unchanged phase values and the trial-shuffled amplitude values. As we randomised the relative trial structure between phase and amplitude while maintaining the temporal structure, and thus left any pulse-evoked changes intact, significant differences to the observed data could not arise from spurious stimulus-evoked relationships between phase and amplitude values [39]. We next averaged each set’s ERPAC data over the electrodes of interest, then took the difference between motor or visual TMS and sham stimulation and averaged resulting values over participants. One observed time-frequency representation and a distribution of 500 surrogate representations emerged, all of which were subsequently binarised by thresholding them with the 95^th^ percentile of the surrogate distribution at each time-frequency point. Contiguous suprathreshold points were clustered and the sum of ERPAC values within each cluster was determined. To account for multiple comparisons, we removed from the observed time-frequency representation those clusters whose cluster sum of ERPAC values was below the 95^th^ percentile of the distribution of maximum cluster sums, obtained by taking the highest sum within each surrogate representation.

Second, to investigate whether the local enhancement of phase-amplitude coupling differed between sTMS and rTMS, we took the mean ERPAC over the local electrodes of interest (C1 and C3 for motor TMS; Oz and O2 for visual TMS), subtracted corresponding mean data obtained from sham stimulation and averaged values over a time window of interest, covering ± 1/10^th^ of the respective phase-providing oscillation’s cycle around either the sTMS pulse or the last pulse of the rTMS trains, as well as over the sixteen amplitude frequencies. By selecting a narrow time window around the last rTMS pulse, we aimed at minimising the potential contamination of the rTMS data from surrounding pulses. Resulting values were then compared between sTMS and rTMS using a two-tailed paired-sample Student’s *t*-test over participants.

Third, we assessed whether an enhancement of phase-amplitude coupling by sTMS and rTMS was observable not only at the stimulation site, but also over other cortical regions. ERPAC was therefore computed at all scalp electrodes for each time point within nine different time windows of interest, centred at −2 cycles to +6 cycles of a condition’s phase-providing oscillation in 1-cycle steps around the onset of the first (or single) pulse and spanning ± 1/10^th^ of this cycle. Topographic maps were created by taking the difference between motor or visual TMS and sham stimulation, and then averaging the resulting values over time points within the respective window of interest, over the sixteen amplitude frequencies as well as over participants.

Fourth, to analyse whether the global propagation of phase-amplitude coupling differed between sTMS and rTMS, we counted the number of electrodes that showed significantly higher ERPAC during motor or visual TMS than during sham stimulation using one-tailed paired-sample Student’s t-tests over participants. Tests were performed for windows of ± 1/10^th^ of a condition’s phase-providing oscillation’s cycle around the sTMS pulse and each of the five rTMS pulses, with ERPAC values averaged over the respective time points as well as over amplitude frequencies. The extent of propagation induced by each of the five rTMS pulses was then compared to the sTMS-induced extent of propagation using exact binomial tests with parameters *n_el_1_* = number of electrodes with a significant TMS-sham difference during a particular rTMS pulse but not the sTMS pulse, *n_el_2_* = number of electrodes with a significant TMS-sham difference during the sTMS pulse but not a particular rTMS pulse and the total number of discordant electrodes *n_el_* = *n_el_1_* + *n_el_2_*. As the assignment of these electrodes to either *n_el_1_* or *n_el_2_* would have happened with equal probability under the null hypothesis of no sTMS-rTMS difference, the *p*-value was defined as the probability of *n_el_1_* reaching the observed or a higher value. Since we performed five tests per condition, multiple comparisons were subsequently accounted for by adjusting *p*-values with the false discovery rate (FDR) procedure [40].

All analyses were performed in MATLAB (The MathWorks, Inc., USA), using the CSD toolbox [34], the CircStat toolbox [38], the FieldTrip toolbox [41] and custom-written scripts.

## 3. Results

### 3.1. Local Modulation of Phase-Amplitude Coupling

Time-frequency representations of the local change in ERPAC relative to the sham stimulation revealed that both motor TMS, analysed at electrodes C1 and C3, and visual TMS, analysed at electrodes Oz and O2, led to an enhancement of phase-amplitude coupling in all assessed phase-amplitude combinations (Figure 2). For sTMS (Figure 2a), significant time-frequency clusters (*p* ≤ .05, one-tailed cluster-based permutation tests) were found around the onset of the single pulse at 0 ms in all conditions but one: The ERPAC increase around the time of the pulse did not reach significance for the effect of visual sTMS on alpha-gamma coupling. However, later clusters of significant increases suggested an effect of sTMS on local ERPAC in this condition as well. For rTMS (Figure 2b), significant time-frequency clusters of increased ERPAC could likewise be observed around the onset times of almost all pulses. Interestingly, whereas the ERPAC increases induced by the individual pulses were clearly separated in time in the 5 Hz and 11 Hz stimulation, which related to theta-gamma and alpha-gamma coupling, respectively, the effects were more strongly merged for the beta-gamma coupling occurring during the faster 23 Hz stimulation. Although clusters in all conditions could spread out symmetrically in time because of the temporal smoothing introduced by the wavelet convolution, it should be noted that their spreading was generally biased towards post-stimulation rather than pre-stimulation time points. While the ERPAC enhancement induced by the present TMS design thus seemed to linger for some tens of milliseconds, it was still transient in nature, with individual effects typically lasting for less than 50 ms. The abovementioned temporal smoothing also explains the observation of enhanced ERPAC during the interpolation intervals, which did not carry meaningful information per se. Since enhanced ERPAC could be found further away from the pulses and interpolation intervals as well (e.g., enhanced theta-gamma coupling more than 3 theta cycles after motor sTMS), these intervals were unlikely to be causally related to the observed effects.

**Figure 2:**
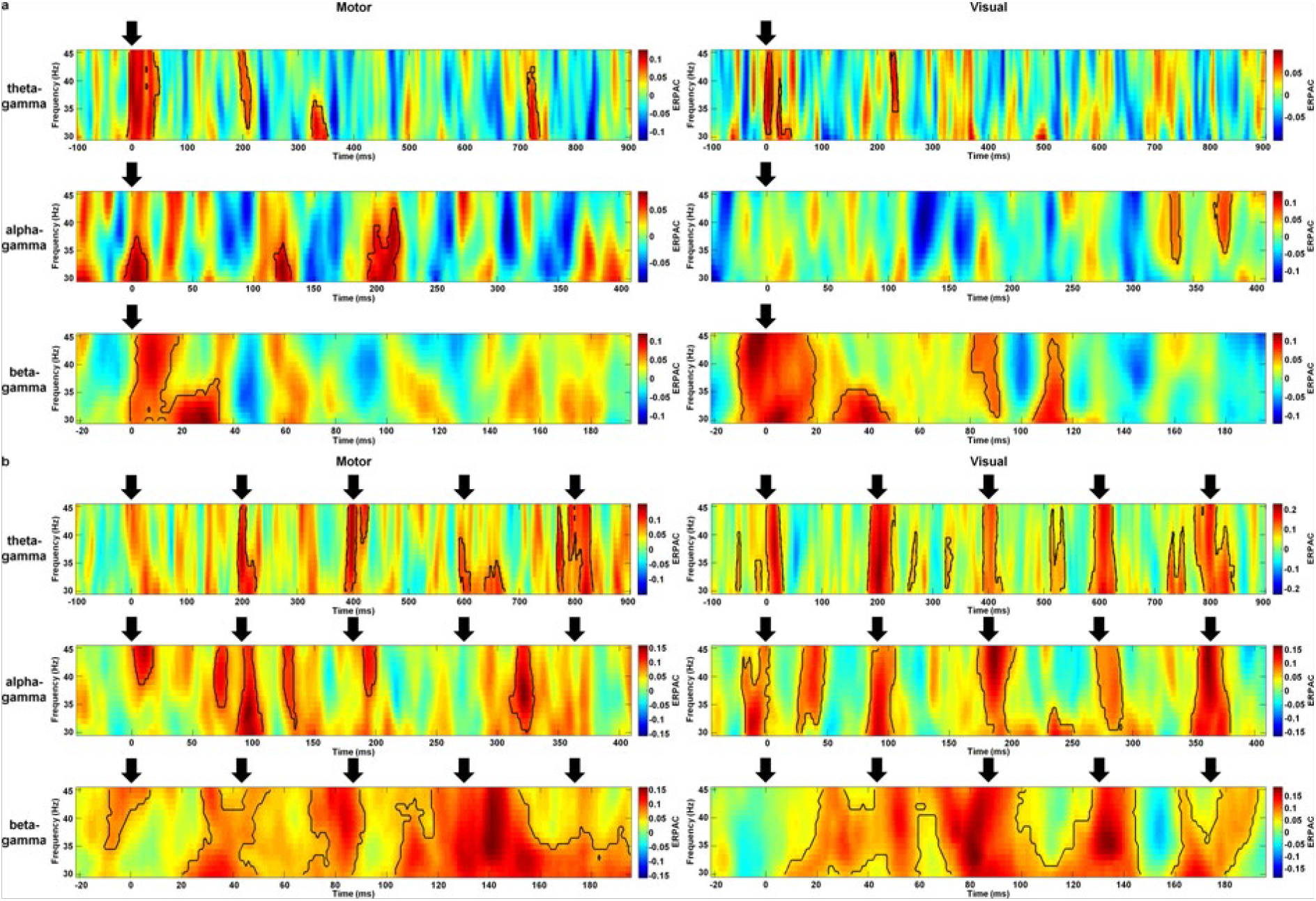
Grand average time-frequency representations of phase-amplitude coupling. Plots show the strength of motor-TMS-induced (left column) and visual-TMS-induced (right column) theta-gamma (first row), alpha-gamma (second row) and beta-gamma (third row) event-related phase-amplitude coupling (ERPAC) as a function of trial time and amplitude frequency. Stimulation paradigms are (a) sTMS and (b) rTMS, with the rTMS frequency always corresponding to the frequency of the phase series. We extracted TMS effects by averaging ERPAC over electrodes C1 and C3 for motor TMS and electrodes Oz and O2 for visual TMS, subtracting corresponding mean data obtained during sham stimulation and averaging the resulting values over the fourteen assessed participants. Time points of pulses are indicated by black arrows and significant time-frequency clusters (*p* ≤ .05, one-tailed cluster-based permutation tests) by black contours.

A comparison of the local change in phase-amplitude coupling induced by sTMS and rTMS revealed that in all but one condition, the mean ERPAC increase relative to the sham stimulation was higher for the last rTMS pulse than for the sTMS pulse, with the opposite pattern being observable for beta-gamma coupling during visual TMS (Figure 3). However, because of high variability over participants, the *p*-values from two-tailed paired-sample Student’s t-tests did not reach statistical significance (all *p* > .05), there being merely a statistical trend (*t*(13) = −1.93, *p* = .076) observable for alpha-gamma coupling during visual TMS, suggesting stronger ERPAC enhancement by rTMS than by sTMS.

**Figure 3:**
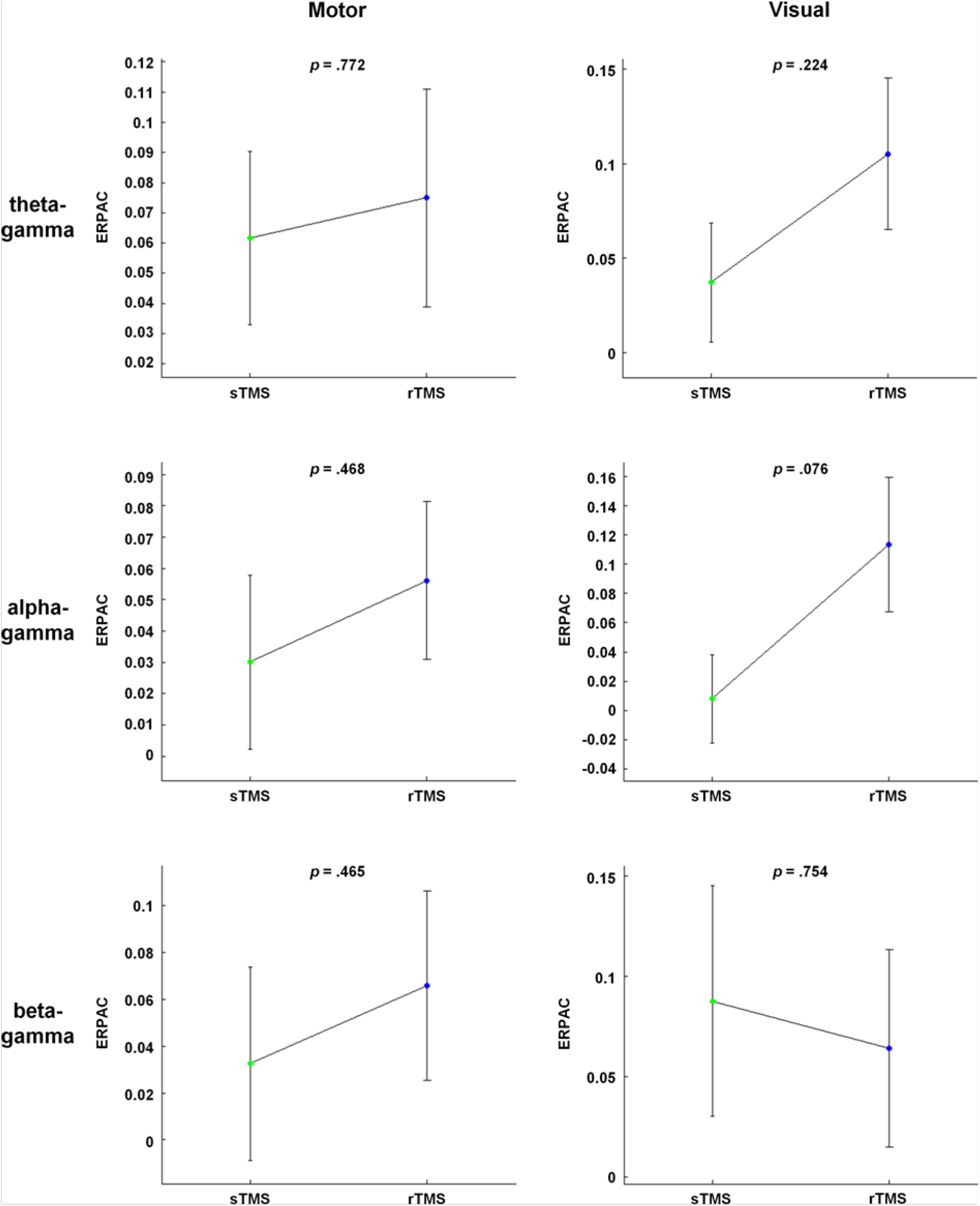
Local comparisons of phase-amplitude coupling between sTMS and rTMS. Plots show the strength of motor-TMS-induced (left column) and visual-TMS-induced (right column) theta-gamma (first row), alpha-gamma (second row) and beta-gamma (third row) event-related phase-amplitude coupling (ERPAC) during the sTMS pulse in green and the last rTMS pulse in blue, with the rTMS frequency always corresponding to the frequency of the phase series. We extracted TMS effects by averaging ERPAC over electrodes C1 and C3 for motor TMS and electrodes Oz and O2 for visual TMS, subtracting corresponding mean data obtained during sham stimulation and averaging the resulting values over predefined time windows of interest around the respective pulses and over the sixteen amplitude frequencies. Bars of sTMS and rTMS represent mean values ± 1 SEM over the fourteen assessed participants; each displayed *p*-value is based on a two-tailed paired-sample Student’s *t*-test between the stimulation paradigms.

### 3.2. Global Modulation of Phase-Amplitude Coupling

To illustrate the change in ERPAC relative to the sham stimulation at all 63 scalp electrodes, nine topographic maps were computed for each condition and stimulation paradigm (Figure 4). The first two maps represented pre-stimulation time windows, the next one (sTMS) or next five (rTMS) represented windows centred on the individual pulses and all remaining maps represented post-stimulation time windows. In accordance with the transient character of the assessed effects, an enhancement of ERPAC was most noticeable within the topographic maps centred on the pulses. Visual inspection further revealed that sTMS-induced increases in ERPAC were prominent primarily over the site of stimulation, with sporadic enhancements also occurring at other sites (Figure 4a). By contrast, the effects of rTMS within the five topographic maps centred on the five pulses appeared to be more strongly distributed over the entire cortex (Figure 4b).

**Figure 4:**
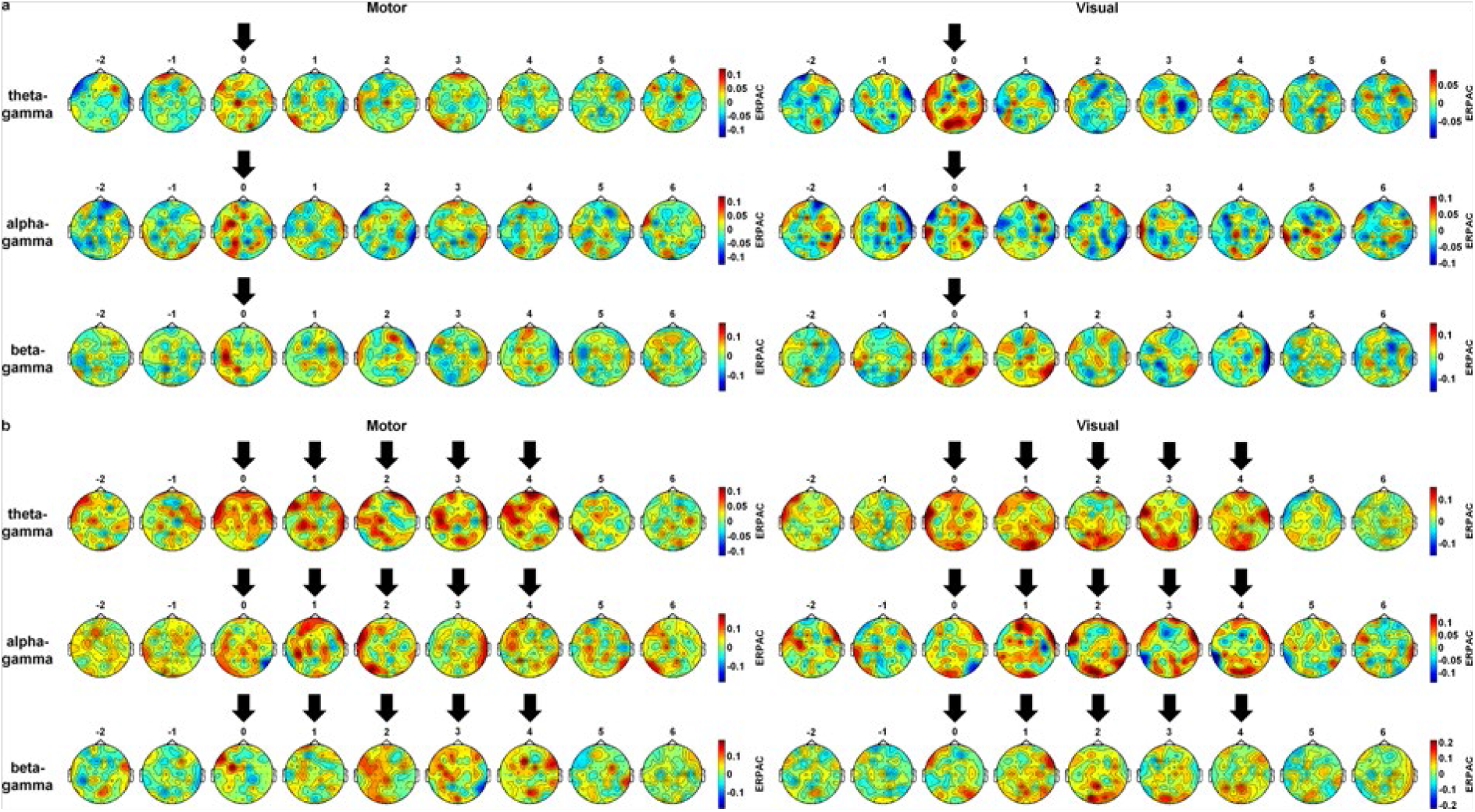
Grand average topographic maps of phase-amplitude coupling. Plots show the strength of motor-TMS-induced (left column) and visual-TMS-induced (right column) theta-gamma (first row), alpha-gamma (second row) and beta-gamma (third row) event-related phase-amplitude coupling (ERPAC) at all scalp electrodes within nine time windows of interest. Stimulation paradigms are (a) sTMS and (b) rTMS, with the rTMS frequency always corresponding to the frequency of the phase series. We extracted TMS effects by subtracting ERPAC obtained during sham stimulation from that obtained during motor or visual TMS and averaging the resulting values over predefined time windows of interest, positioned at −2 cycles to +6 cycles of a condition’s phase-providing oscillation around the onset of the first (or single) pulse, over the sixteen amplitude frequencies and fourteen assessed participants. Topographic maps centred on pulses are indicated by black arrows.

We quantified this observation by determining the number of electrodes with significantly higher ERPAC during motor or visual TMS than during sham stimulation (*p* ≤ .05, one-tailed paired-sample Student’s *t*-tests), and then comparing the electrode numbers between sTMS and the five rTMS pulses (Figure 5). As expected, in most cases, the number of significant electrodes was larger for a particular rTMS pulse than for the sTMS pulse of the same condition. With regard to motor stimulation, this difference was statistically significant (*p_FDR_* ≤ .05, exact binomial tests) for three out of five rTMS pulses when investigating alpha-gamma coupling (pulses 1, 2, 3: each *p_FDR_* = .041) and for one rTMS pulse when investigating beta-gamma coupling (pulse 3: *p_FDR_* = .019). With regard to visual stimulation, three out of five rTMS pulses showed a significantly larger propagation when investigating theta-gamma coupling (pulse 1: *p_FDR_* = .019, pulse 4: *p_FDR_* = .009, pulse 5: *p_FDR_* = .006), whereas two rTMS pulses were significant for beta-gamma coupling (pulses 2, 3: each *p_FDR_* = .021). Thus, while significant sTMS-sham differences were still found at five or more electrodes in all conditions, indicating a certain extent of propagation in this stimulation paradigm as well, rTMS induced a considerably more widespread propagation of ERPAC enhancement overall.

**Figure 5:**
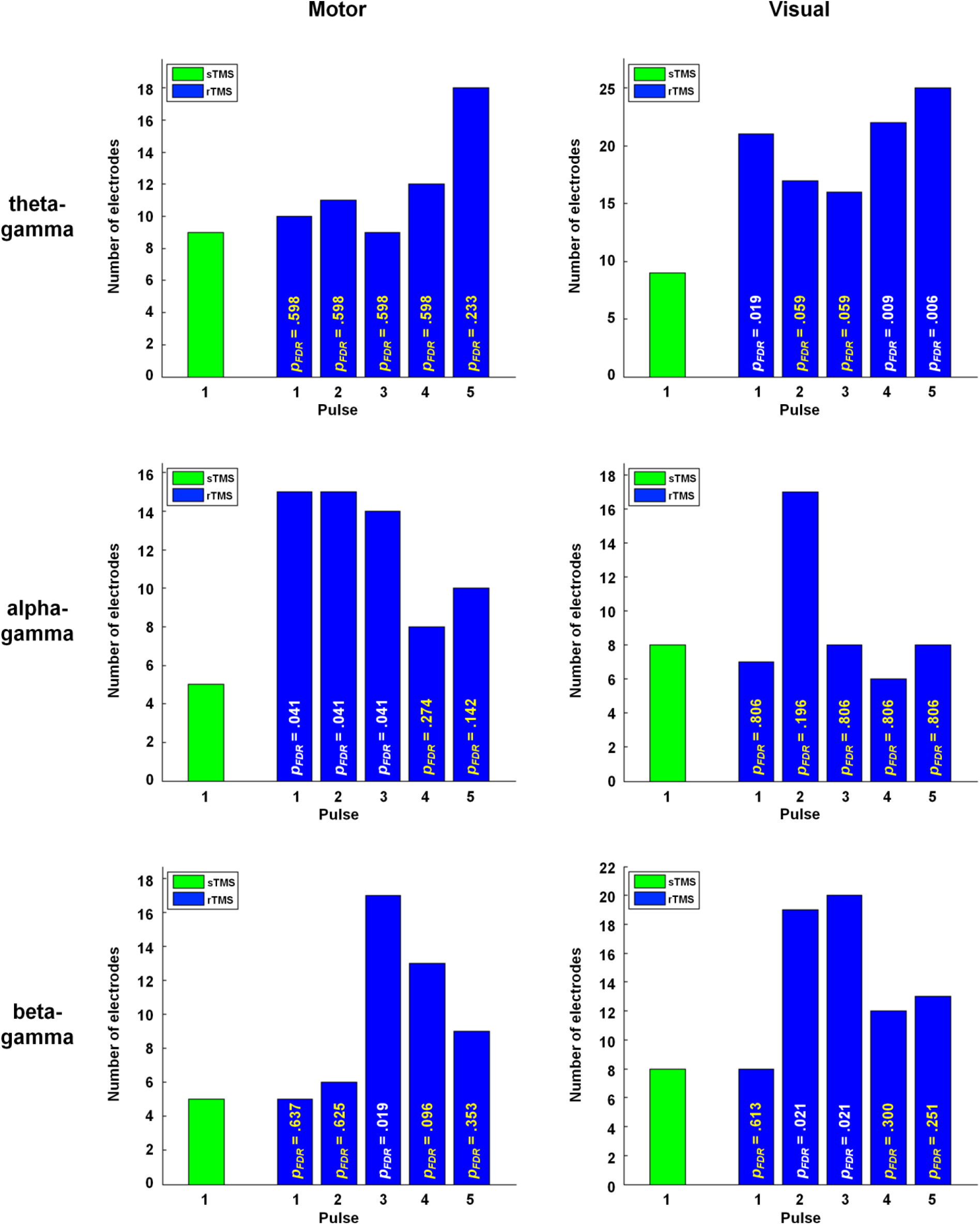
Global comparisons of phase-amplitude coupling between sTMS and rTMS. Plots show the spatial extent of motor-TMS-induced (left column) and visual-TMS-induced (right column) theta-gamma (first row), alpha-gamma (second row) and beta-gamma (third row) event-related phase-amplitude coupling (ERPAC) during the sTMS pulse in green and the five rTMS pulses in blue, with the rTMS frequency always corresponding to the frequency of the phase series. We extracted TMS effects by averaging ERPAC at all electrodes over predefined time windows of interest around the respective pulses and over the sixteen amplitude frequencies, comparing the resulting values between motor or visual TMS and sham stimulation with one-tailed paired-sample Student’s t-tests and counting the number of electrodes with a statistically significant difference (*p* ≤ .05; drawn on y-axis). The spatial extent of TMS effects was subsequently compared between the sTMS pulse and each of the five rTMS pulses with exact binomial tests. Corresponding *p*-values, adjusted for multiple comparisons with the false discovery rate (FDR) procedure, are superimposed onto the rTMS bars in all plots; significant *p*-values (*p_FDR_* ≤ .05) are displayed in white, all other *p*-values in yellow.

## 4. Discussion

With the present study, we provide compelling evidence that both sTMS and rTMS can transiently enhance phase-amplitude coupling of neural oscillations, as measured with concurrent EEG. This enhancement was found not only locally at the stimulation site, but also over various other cortical sites, with the propagation induced by rTMS outperforming that induced by sTMS. By demonstrating enhanced theta-gamma, alpha-gamma and beta-gamma phase-amplitude coupling during motor and visual TMS, our results have relevance for a wide range of the nested oscillatory signatures inherent to neural processing [20, 21, 42, 43] and are highly consistent with the hypothesised population-level increase in intrinsic coupling brought about by oscillatory phase alignment. We hence propose that concurrent TMS-EEG can be utilised to effectively probe such coupling in humans, a feature making it a highly promising technique for future non-invasive investigations of this important mechanism.

At the site of stimulation, all of the assessed conditions showed significant increases in phase-amplitude coupling strength during or slightly after TMS. As the phase-amplitude coupling in the present study was operationalised as the circular-linear correlation of phase and amplitude values at each time point across stimulation trials [38, 39], changes in coupling strength could be assessed without the loss of temporal resolution inherent to most other coupling measures [21, 22, 29]. Given that the enhancement of local phase-amplitude coupling typically lasted for less than 50 ms around the pulse, a finding consistent with the previously reported short-lived character of TMS-induced phase dynamics [13], this approach was vital to quantify transient effects that would otherwise be barely detectable in scalp EEG recordings. We took the following steps to ensure that the observed effects did indeed reflect a direct enhancement of macroscopic phase-amplitude coupling by TMS. First, to account for any indirect effects of stimulation, particularly for auditory-evoked changes in brain activity including cross-modally triggered phase locking after salient sounds [44], phase-amplitude coupling was always assessed relative to the sham stimulation, which was applied over the vertex of the head. Second, by statistically comparing the observed TMS-sham differences to surrogate distributions of trial-shuffled data with unmodified temporal structure [39], we confirmed that the observed enhancement of phase-amplitude coupling was based on a specific statistical relationship between phase and amplitude values across trials, rather than on spurious relationships induced by unrelated neural effects of the pulse or any sharp edge artefacts [45]. Thanks to these methodological approaches, a clear demonstration of TMS-induced changes in phase-amplitude coupling was made possible. Even though such changes seemed to be stronger for the last pulse of the delivered rTMS trains (targeted at the phase-providing lower-frequency oscillations) than for the sTMS pulse in almost all conditions, the statistical power was not high enough to enable a conclusion regarding local phase-amplitude coupling differences between stimulation paradigms. It has to be acknowledged in this context that 30 collected trials per block might have been insufficient to yield significant difference effects. Due to the already long overall testing duration of 4–5 h per participant though, a higher number of trials was practically not feasible in our study. Notably, a slight modification of the rTMS paradigm could potentially facilitate the detection of significant local differences. Successful neural entrainment, which might underlie a potential rTMS benefit by enabling stronger oscillatory phase alignment relative to sTMS, requires the existence of a neural population that can oscillate at the stimulation frequency under natural conditions [17]. As such eigenfrequencies differ between cortical regions [15] and individuals [46], the entrainment capability of rTMS should be enhanced by tuning its frequency to the local power spectrum peak frequencies of participants. Recent evidence has indeed demonstrated the benefits of such an individualised targeting of intrinsic oscillations by rTMS [14], making a comparison of local phase-amplitude coupling strength between this rTMS paradigm and sTMS promising. Importantly, even though the perturbation of intrinsic oscillations is potentially stronger in the case of individualised rTMS frequencies, previously reported effects of non-individualised stimulation on human cognition [47] suggest successful entrainment in this case as well. As Thut and colleagues [12] noted, such effects might be enabled by intra-individual frequency fluctuations as well as a loosening relationship between eigenfrequency and effective stimulation frequency at higher stimulation intensities (see also Gouwens and colleagues [48]). A decision in favour of non-individualised stimulation paradigms may eventually also be driven by the increased expenditure of time and resources associated with the pre-experimental determination of individual peak frequencies, especially when testing clinical populations.

Alongside the described local effects of sTMS and rTMS, we found that TMS can enhance population-level measures of phase-amplitude coupling over various other cortical sites. In line with this finding, propagation of neural activation related to either sTMS or rTMS has been shown in a number of previous studies [13, 14, 49–54]. In particular, Kawasaki and colleagues [13] demonstrated an sTMS-induced large-scale propagation of oscillatory phase locking, which was accompanied by increased directional information flow of phase dynamics from the occipital stimulation site to an examined distant site over the motor cortex, as assessed by transfer entropy. Since the therein suggested alignment of phases of individual oscillators should increase the detectability of intrinsic phase-amplitude coupling at the population level, the propagation of enhanced coupling observed here is highly consistent with this report. The pathways of such propagation are not arbitrary, but should follow the brain’s intrinsic organizational structure, which is characterized by frequency-specific functional networks [55]. We thus propose that the applied stimulation drove particular cortical systems via successive interactions with functionally coupled neural oscillators in a frequency-dependent manner, resulting in different propagation patterns for the different conditions. However, knowledge of any propagation differences between sTMS and rTMS is sparse. In the present study, rTMS enhanced scalp-recorded phase-amplitude coupling at considerably more sites than sTMS. This difference was particularly pronounced for alpha-gamma coupling during 11 Hz motor stimulation and theta-gamma coupling during 5 Hz visual stimulation, with three out of five rTMS pulses outperforming the respective sTMS pulse in each case. As (nested) intrinsic oscillations are believed to play an important role in neural signal transmission [42, 56], their entrainment by rTMS may again be at the bottom of the observed benefit. In accordance with this idea, Romei and colleagues [14] demonstrated that rTMS pulses propagate from the sensorimotor cortex to spinal levels only when sensorimotor oscillations are specifically targeted via their eigenfrequency, with stimulation at other frequencies having little impact on cortico-spinal signal interactions. Likewise, the impact of sTMS on relevant oscillations might have been too weak to reach the extent of propagation achieved by rTMS in the present study. Before alternatively ascribing the observed propagation benefit of rTMS to a methodological contamination of rTMS pulses by surrounding pulses, it should be noted that during rTMS, both local alpha-gamma coupling and local theta-gamma coupling typically returned to baseline long before the next pulse arrived. Still, one could argue that we had already observed a more widespread distribution of enhanced phase-amplitude coupling during the first rTMS pulse in several conditions. As an rTMS-induced synchronisation of neural oscillators might have progressively strengthened within entire rTMS blocks, including multiple stimulation trials, in these cases, the assessed correlation of phase and amplitude values could have been driven by the intensified entrainment present only in later trials. Nonetheless, conclusive evidence on this matter is so far missing and future studies are needed to shed light on the exact cause of the observed sTMS-rTMS differences in phase-amplitude coupling propagation.

As phase-amplitude coupling is believed to play a fundamental role in the transfer of neural information across diverse spatial and temporal processing scales, thereby serving the dynamic integration of global computations with fast local processing, it may be extremely relevant for cognitive functioning [42]. Recent evidence has started to support this claim by hinting at its functional significance for visual perception [26], feedback processing [57], memory recall [58], visuomotor mapping [59] and movement planning and execution [60]. Accordingly, a dysfunction in phase-amplitude coupling has been identified in several clinical conditions such as Parkinson’s disease [61], autism spectrum disorders [62] and epilepsy [63]. Still, our current understanding of this intriguing mechanism is far from exhaustive. Investigations of phase-amplitude coupling in the human population are hampered by the inherent shortcomings of established non-invasive measurement techniques. As methods such as EEG capture the summed potentials of tens of thousands of synchronously activated neurons, scalp-recorded oscillations inevitably reflect the summation of multiple underlying neural oscillators. Consequently, even strong phase-amplitude coupling can only be detected with EEG if a considerable quantity of those oscillators are in phase, and thus are not cancelling out at the population level. We suggest that by aligning the phases of individual oscillators, TMS fosters this setting, and thereby facilitates the non-invasive detection of intrinsic phase-amplitude coupling with an improved signal-to-noise ratio. The proposed perturbational approach therefore holds great promise for future investigations aimed at further unravelling the association between such oscillatory nesting on the one hand, and healthy or pathological human functioning on the other hand. By probing the intrinsic capacity of individuals for phase-amplitude coupling, concurrent TMS-EEG might, in this regard, prove particularly useful for the reliable development of coupling-based biomarkers, as have already been presented for amnestic mild cognitive impairment [64]. Besides opening the door to a deeper understanding of the functional role of phase-amplitude coupling, the present results add to the constantly growing body of knowledge regarding the neurophysiological mode of action underlying TMS (see Klomjai and colleagues [65] for a review). By actively modulating nested intrinsic oscillations, TMS impacts on the gating of information along interconnected neural ensembles, and consequently affects a fundamental property of neural processing in the human brain.

On a final note, we would like to point out that direct evidence for this assumed mode of action is still missing. Even though our results are highly consistent with the hypothesised population-level increase in intrinsic phase-amplitude coupling, it cannot be fully excluded that TMS instead gave rise to novel coupling overlaid on top of ongoing brain activity (see Sauseng and colleagues [16] for a similar discussion on the generation of event-related potentials). One might argue in favour of the latter mechanism in particular by pointing at the lack of observable differences between motor and visual TMS in our study. However, the above-mentioned loosened relationship (at higher stimulation intensities) between a region’s eigenfrequency, which most likely differs between motor and visual cortex [15], and the region’s effective stimulation frequency may have contributed to this observation as well. The former, hypothesised mechanism on the contrary might be supported by evidence for the pre-TMS existence of relevant phase-amplitude coupling that is then enhanced after TMS. While the current ERPAC method is not suited for the assessment of non-event-related data, phase-amplitude coupling has indeed been demonstrated in the human resting state [23, 27]. The presence or absence of coupled macroscopic oscillations alone, however, is an insufficient marker of the existence of underlying intrinsic oscillators [16], which in turn does not automatically entail that these oscillators are directly modulated by TMS. Thus, although TMS has generally been shown to interact with intrinsic brain activity [66], the unequivocal disentanglement of both mechanisms in the context of the present study requires access to the level of individual oscillators with measurements of a considerably higher temporal and spatial resolution compared to that achieved by time-frequency-resolved scalp EEG.

In conclusion, we used a concurrent TMS-EEG study design to demonstrate that TMS can transiently enhance scalp-recorded phase-amplitude coupling. This enhancement was found for both sTMS and rTMS, with a more widespread propagation of effects being observed during the latter stimulation paradigm. We thus recommend the perturbational approach of concurrent TMS-EEG as a novel experimental technique to effectively probe intrinsic phase-amplitude coupling in humans. The utility of this design for future studies investigating the functional roles of phase-amplitude coupling in the healthy population, as well as plastic changes of phase-amplitude coupling in pathological conditions, awaits confirmation.

## Data Availability

The TMS-EEG data used to support the findings of this study are available from the corresponding author upon request.

